# MAVS polymers smaller than 80 nm induce mitochondrial membrane remodeling and interferon signaling

**DOI:** 10.1101/506568

**Authors:** Ming-Shih Hwang, Jérôme Boulanger, Jonathan Howe, Anna Albecka, Mathias Pasche, Leila Mureşan, Yorgo Modis

**Affiliations:** Department of Medicine, University of Cambridge, MRC Laboratory of Molecular Biology, Francis Crick Avenue, Cambridge, CB2 0QH, United Kingdom; MRC Laboratory of Molecular Biology, Francis Crick Avenue, Cambridge, CB2 0QH, United Kingdom; Cambridge Advanced Imaging Centre, University of Cambridge, Cambridge CB2 1QP, United Kingdom; Department of Physiology, Development and Neuroscience, University of Cambridge, Cambridge CB2 1QP, United Kingdom

**Keywords:** innate immunity, pathogen-associated molecular pattern (PAMP), signal transduction, STORM, cell death

## Abstract

Double-stranded RNA (dsRNA) is a potent proinflammatory signature of viral infection. Oligomerization of RIG-I-like receptors on cytosolic dsRNA nucleates self-assembly of the mitochondrial antiviral signaling protein (MAVS). In the current signaling model, the caspase recruitment domains of MAVS form helical fibrils that self-propagate like prions to promote signaling complex assembly. However, there is no conclusive evidence that MAVS forms fibrils in cells or with the transmembrane anchor present. We show here with super-resolution light microscopy that MAVS activation by dsRNA induces mitochondrial membrane remodeling. Quantitative image analysis at imaging resolutions as high as 32 nm shows that in the cellular context MAVS signaling complexes and the fibrils within them are smaller than 80 nm. The transmembrane domain of MAVS is required for its membrane remodeling, interferon signaling and proapoptotic activities. We conclude that membrane tethering of MAVS restrains its polymerization and contributes to mitochondrial remodeling and apoptosis upon dsRNA sensing.

## Introduction

Recognition of viral nucleic acids by innate immune receptors is one of the most conserved and important mechanisms for sensing viral infection. Many viruses deliver or generate doublestranded RNA (dsRNA) in the cytosol of the host cell. Cytosolic dsRNA is a potent proinflammatory signal in vertebrates. Endogenous dsRNAs are modified or masked through various mechanisms to prevent autoimmune signaling, and genetic deficiencies in these dsRNA modification pathways can cause autoimmune disorders [1–3]. Cytosolic dsRNA is primarily sensed by the RIG-I-like receptors (RLRs) RIG-I (*DDX58*), MDA5 (*IFIH1*) and LGP2 (*DHX58*) [4], which activate the mitochondrial antiviral signaling protein (MAVS) [5–8]. RIG-I recognizes dsRNA blunt ends with unmethylated 5’-di- or triphosphate caps [9–12]. MDA5 recognizes uninterrupted RNA duplexes longer than a few hundred base pairs [11, 13]. LGP2 functions as a cofactor for MDA5 by promoting the nucleation of MDA5 signaling complexes near dsRNA blunt ends [14, 15]. Binding to dsRNA causes RIG-I to form tetramers and MDA5 to cooperatively assemble into helical filaments around the dsRNA [16–19]. RIG-I and MDA5 each contain two N-terminal caspase recruitment domains (CARDs). The increased proximity of the CARDs upon RLR oligomerization induces the CARDs from four to eight adjacent RLR molecules to form a helical lock-washer-like assembly [19, 20]. These helical RLR CARD oligomers bind to MAVS, which has a single N-terminal CARD, via CARD-CARD interactions [5]. Binding of MAVS CARDs to RLR CARD oligomers nucleates the polymerization of MAVS CARD fibrils with amyloid-like (or prion-like) properties including resistance to detergents and proteases [19–21]. MAVS polymerization is required for signaling, and the spontaneous elongation of MAVS fibrils following nucleation is thought to provide a signal amplification mechanism [21]. MAVS fibrils then recruit proteins from the TRAF and TRIM families to form multimeric signaling platforms, or signalosomes [21]. MAVS is localized primarily on the outer mitochondrial membrane [5] but can also migrate via the mitochondria-associated membrane (MAM) to peroxisomes [22], which function as an alternative signaling platform to mitochondria [23]. MAVS signalosomes activate both type I interferon (through IRF3) and NF-ĸB-dependent inflammatory responses [11, 13, 21]. Overexpression of MAVS induces apoptotic cell death, and this proapoptotic activity is dependent on its transmembrane anchor (TM) and mitochondrial localization, but independent of the CARD [24, 25]. A loss-of-function MAVS variant is associated with a subset of systemic lupus patients [26].

In the current model of MAVS signaling, RLR CARD oligomers trigger a change of state in the CARD of MAVS, from monomer to polymeric helical fibril. MAVS fibrils grow like amyloid fibrils by drawing in any proximal monomeric MAVS CARDs [20]. This model is based partly on the observation that purified monomeric MAVS CARD spontaneously assembles into fibrils of 0.2 - 1 μm in length [19, 21]. The fibrils, but not the monomers, activate IRF3 in signaling assays [21] with cell-free cytosolic extracts. Moreover, a purified MAVS fragment lacking the TM can, in its polymeric fibril form, activate IRF3 in crude mitochondrial cell extracts that contain endogenous wild-type MAVS (17). However, this signaling model is based primarily on signaling assays and structural studies performed in a cell-free environment with soluble fragments of MAVS lacking the TM. MAVS has been reported to form rod-shaped puncta on the outer mitochondria membrane upon activation with Sendai virus [27], but evidence that MAVS forms polymeric fibrils in cells remains inconclusive, and furthermore MAVS fibrils are not sufficient for signaling. Indeed, the MAVS TM is required for interferon induction and cell death activation [5, 21, 25], and several viruses including hepatitis C virus suppress type I interferon production by cleaving it off [8, 28–31]. The sequence between the CARD and TM of MAVS, which represents 80% of the MAVS sequence, is also required for downstream signaling [32]. How this sequence and the TM function together with the CARD in cell signaling remains unclear.

In the cellular context, MAVS CARD fibrils are subject to multiple physical constraints, including tethering to RLR-dsRNA complexes (at one end of the fibril) and to the mitochondrial membrane (via the TM of each MAVS molecule in the fibril). Here, we address the question of how the current model of MAVS signaling can be reconciled with these physical constraints, and the requirement of the TM, for cell signaling. Imaging of MAVS signaling complexes by super-resolution light microscopy with effective optical resolutions of up to 32 nm reveal that in the cellular context MAVS signaling complexes are significantly smaller than expected—no more than 80 nm. Moreover, MAVS signaling is associated with remodeling of mitochondrial compartments and apoptosis, and both of these activities are dependent on the TM of MAVS. Our data indicate that MAVS forms smaller signaling complexes than previously thought [21, 27].

## Results

### MAVS activation by cytosolic RNA induces mitochondrial membrane remodeling

The polymerization of purified soluble fragments of MAVS into helical fibrils is well documented [19, 21]. In the cellular context, however, MAVS is tethered to the outer mitochondrial membrane via its transmembrane anchor, and binds via its CARD to oligomeric or polymeric RLR-dsRNA complexes [19, 21, 33]. To examine how the physical constraints imposed by membrane tethering and association with RIG-I or MDA5 may affect MAVS CARD fibril formation, we imaged cells containing active MAVS signaling complexes by super-resolution fluorescence microscopy. Mouse embryonic fibroblasts (MEFs) and 3T3 cells were imaged by structured illumination microscopy (SIM) and stimulated emission depletion microscopy (STED). Because fluorescent proteins fused to the N-terminus, C-terminus or juxtamembrane region of MAVS were not suitable for STORM, cells were labeled with a monoclonal antibody against a linear epitope within residues 1-300 of MAVS and a fluorescently-labeled secondary antibody. An antibody with an overlapping epitope was shown previously to recognize MAVS in the fibril form in non-reducing semi-denaturing electrophoresis [21]. Immunofluorescence of MAVS and TOM20, an outer mitochondrial membrane marker, showed that the two proteins localized to the same mitochondrial compartments (Fig. 1). We observed changes in the distribution of MAVS and in overall mitochondrial morphology (using the TOM20 marker) upon infection of 3T3 cells with a West Nile virus (WNV) replicon (Fig. 1A). Infection with the WNV replicon caused mitochondrial compartments to form more fragmented and less filamentous structures closer to the nucleus. Introducing the dsRNA mimic poly(I:C) into MEFs by electroporation (0.6 - 1 picogram per cell) recapitulated the changes in MAVS distribution and mitochondrial morphology observed upon infection with the WNV replicon (Fig. 1B). These changes were also recapitulated by cotransfecting MEFs derived from MAVS knockout mice (MAVS KO MEFs) [34] with poly(I:C) RNA and a plasmid encoding MAVS (0.6 - 1 pg of each per cell, Figs. 1C). Mitochondrial remodeling associated with poly(I:C) treatment was quantified as statistically significant reductions in: (1) the distance from the nucleus (from an average of 9.0 μm to 4.5 μm, *p* = 2.0 x 10^-4^) (Fig. 1E); (2) the fraction of the cytosolic area occupied by mitochondria (from an average of 24.2% to 16.7%, *p* = 0.0015) (Fig. 1F); and (3) the length of mitochondrial compartments measured as unbranched segments of skeletonized T0M20 fluorescence (from an average of 2.42 μm to 1.98 μm, *p* = 0.0478) (Fig. 1G). The amount of MAVS plasmid and the electroporation method used in transfections were selected to yield MAVS expression levels that were in the physiological range (see below). Notably, no MAVS filaments longer than the resolution limit were observed. The resolution of SIM and STED was approximately 110 nm and 80 nm in the imaging (*xy*) plane, respectively (and 350 and 600 nm in *z*, respectively). These resolutions were also sufficient to resolve differences in the positions of individual MAVS and T0M20 protein complexes so that the immunofluorescence signals from the two proteins formed an alternating pattern within mitochondrial compartments rather than strictly colocalizing (Figs. 1A, 1C, 2A). We confirmed that transfection with poly(I:C) RNA induced translocation of IRF3 to the nucleus, which is a hallmark of interferon-β (IFN-β) signaling (Fig. 2B). Importantly, we showed that the level of MAVS expression in transfected MAVS KO MEFs was comparable to the physiological level of endogenous MAVS expression in wild-type MEFs (Fig. 3). A live-cell dual-luciferase reporter assay was used to confirm that IFN-β signaling is activated in MAVS KO MEFs transfected with the MAVS expression plasmid and poly(I:C) RNA (see below). Costaining for MAVS and MDA5 (Fig. 2C) showed an increased interaction between the two proteins, defined as the average distance between MAVS and MDA5 fluorescence (Fig. 2D) [35]. However, MDA5 staining remained predominantly cytosolic and no significant increase colocalization was detected with poly(I:C) treatment based on the Pearson correlation. Similarly, RIG-I was recently shown to partition into a MAVS-associated mitochondrial fraction and a cytosolic stress granule fraction [36]. In the absence of MAVS, mitochondria retained their filamentous morphology and failed to move towards the nucleus upon transfection with poly(I:C) (Figs. 1D). We note that the immunofluorescence and cell signaling data show relatively high levels of mitochondrial remodeling (Fig. 1), IRF3 nuclear translocation (Fig. 2B) and background signaling (see below) in cells transfected with a control plasmid instead of poly(I:C), which is likely attributable to IFN-β signal transduction by cytosolic DNA sensors. We conclude that activation of MAVS by cytosolic RNA sensing is associated with remodeling of mitochondria into more globular and perinuclear compartments. Mitochondrial remodeling from the healthy filamentous morphology to perinuclear globular compartments through mitochondrial fission events is a hallmark of apoptosis [37]. Indeed, MAVS was shown previously to promote apoptosis independently of its function in initiating interferon and NF-ĸB signaling [24].

**Fig. 1.**
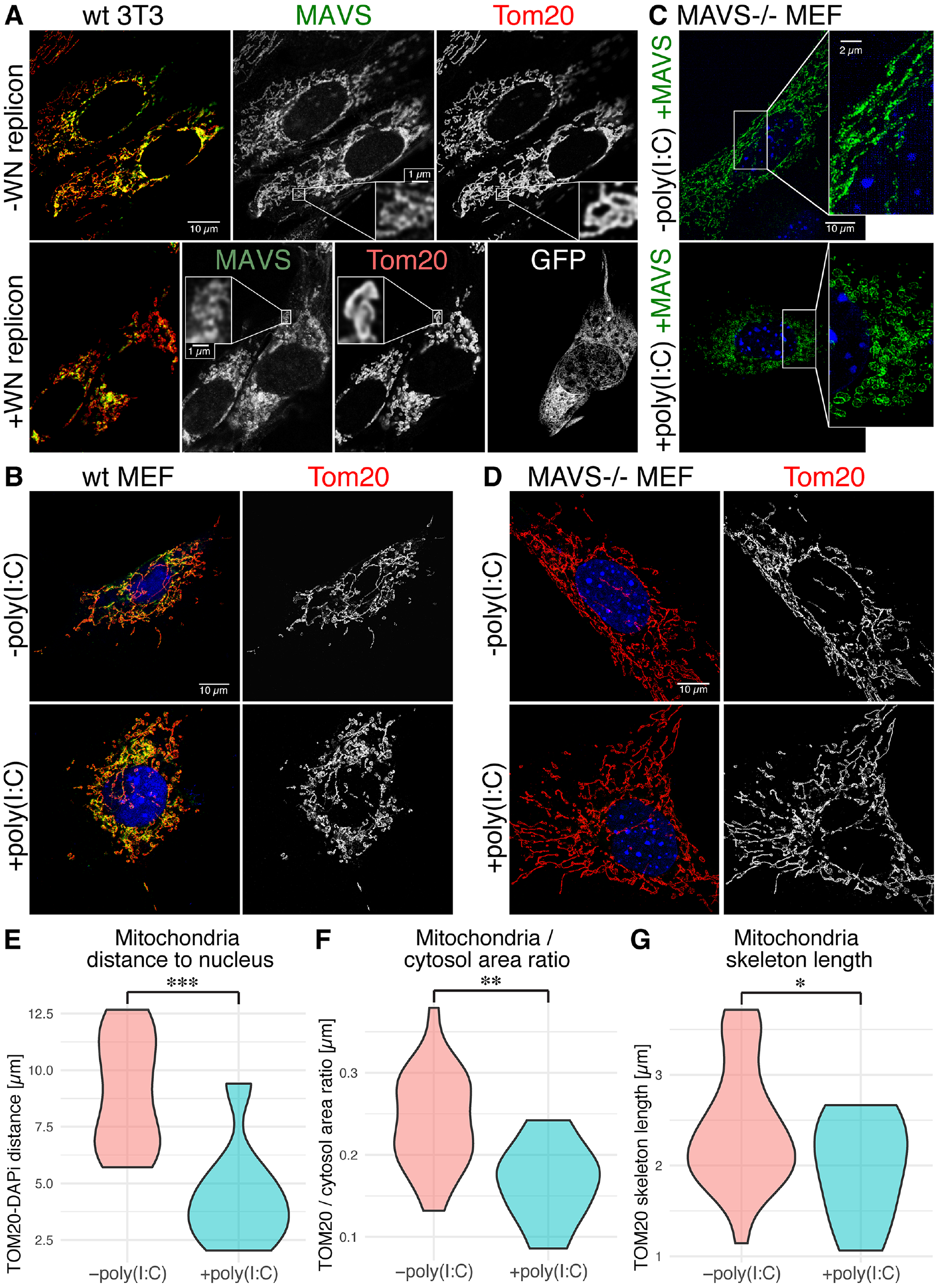
Super-resolution and confocal fluorescence microscopy of MAVS signaling complexes on the outer mitochondrial membrane upon activation with poly(I:C) RNA or a West Nile reporter virus. (**A**) Stimulated emission depletion microscopy (STED) of MAVS and TOM20 in NIH 3T3 cells infected with a GFP-labeled West Nile reporter virus (WN replicon), with immunofluorescently labeled MAVS in green and TOM20 immunofluorescence in red, or with the MAVS staining, TOM20 staining or the GFP infection marker shown separately in grey. Yellow results from green-red overlap and is indicative of MAVS-TOM20 colocalization. (**B**) Confocal images of MAVS and TOM20 in wild-type MEFs, with MAVS (green) and TOM20 (red) immunofluorescence, or with TOM20 staining shown separately in grey. Cells were transfected with either poly(I:C) RNA (+poly(I:C)) or an empty plasmid (-poly(I:C)) to control for the effects of transfection. (**C**) Structured illumination microscopy (SIM) of MAVS immunofluorescence (green) in MAVS KO MEFs cotransfected with MAVS and poly(I:C) RNA (+poly(I:C)), or MAVS and a control plasmid (-poly(I:C)). (**D**) Confocal images of TOM20 in MAVS KO MEFs transfected with either poly(I:C) RNA (+poly(I:C)) or an empty plasmid (-poly(I:C)) but no plasmid encoding MAVS, with immunolabeling of MAVS (green) and TOM20 (red), or with TOM20 staining shown separately in grey. DAPI nuclear staining is shown in blue (panels **B-D**). (**E**) Average distance of TOM20 fluorescence from the nucleus, defined as DAPI fluorescence. (**F**) Quantification of the fraction of the cytosolic area occupied by TOM20 fluorescence, from bright field immunofluorescence imaging (100x magnification) of 39 cells (28 −poly(I:C), 11 +poly(I:C)). (**G**) Average length of mitochondrial compartments measured as the length of unbranched segments of skeletonized TOM20 fluorescence, from the same images used in (**F**).

**Fig. 2.**
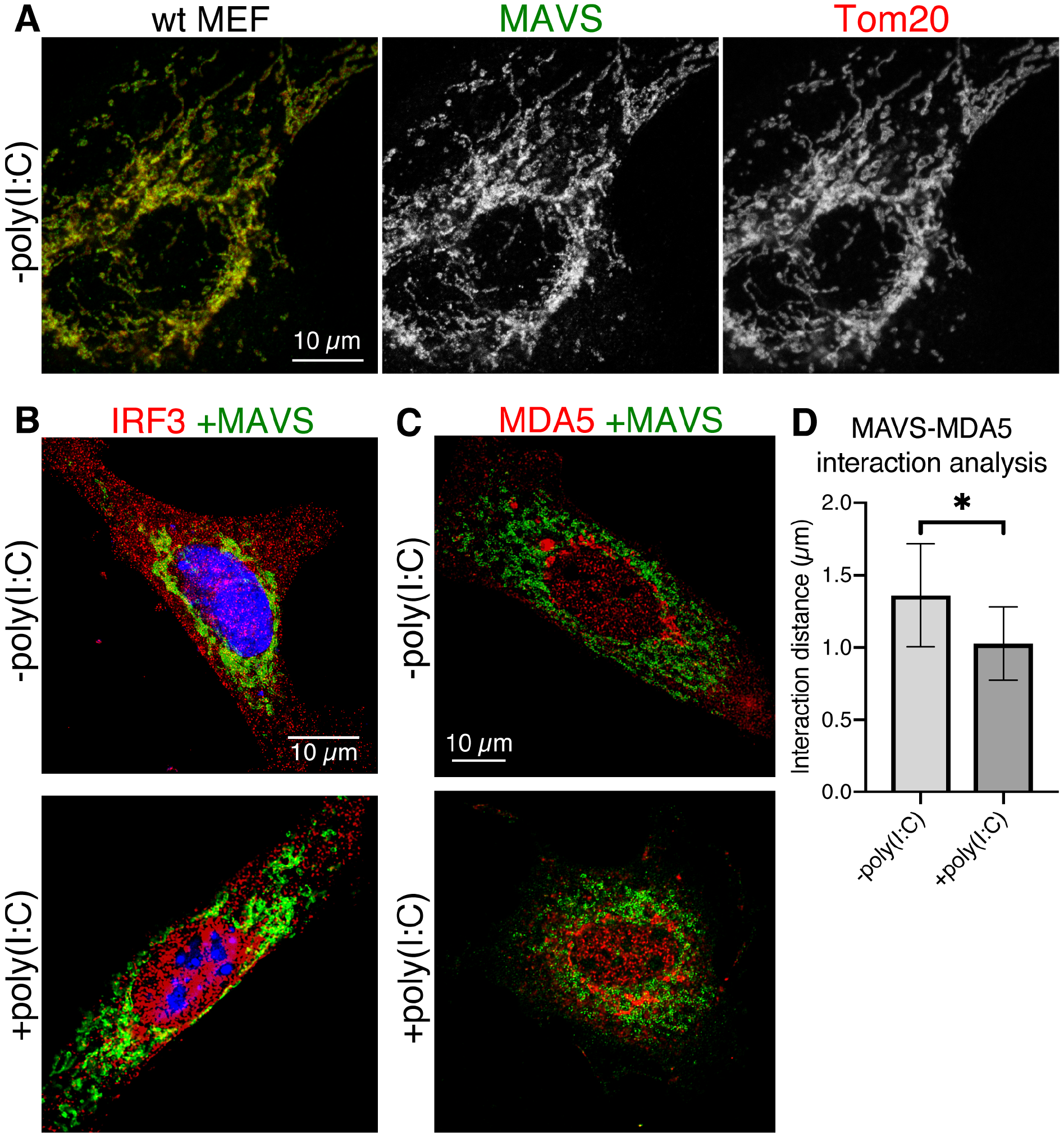
Immunofluorescence microscopy showing mitochondrial morphology and cellular localization of IRF3 and MDA5 upon MAVS activation with poly(I:C) RNA. (**A**) Baseline mitochondrial morphology of wild-type MEFs without any transfection of plasmid DNA or poly(I:C) RNA. Immunolabeled MAVS and TOM20 are in green and red, respectively (left), and separately in grey (center and right). (**B**) Nuclear translocation of IRF3 on MAVS activation with poly(I:C) RNA. Representative image of MAVS KO MEFs cotransfected with either wild-type MAVS and a control plasmid (-poly(I:C)) or with wild-type MAVS and poly(I:C) RNA (+poly(I:C)). MAVS and IRF3 immunofluorescence signals are green and red, respectively. DAPI nuclear staining is blue. (**C**) Representative image of MAVS KO MEFs transfected either with wild-type MAVS and a control plasmid (-poly(I:C)) or with wild-type MAVS and poly(I:C) RNA (+poly(I:C)). MAVS and IRF3 signals are green and red, respectively. (**D**) Interaction analysis of MAVS and MDA5 fluorescence. The average distance between MDA5 and MAVS points was 32% smaller in cells transfected with MAVS (1.03 μm) versus cells transfected with control plasmid DNA (1.36 μm). Error bars represent the standard deviation from the mean; n = 4. Statistical significance (p = 0.019) was calculated in Prism 8 with a 1-sided t-test.

**Fig. 3.**
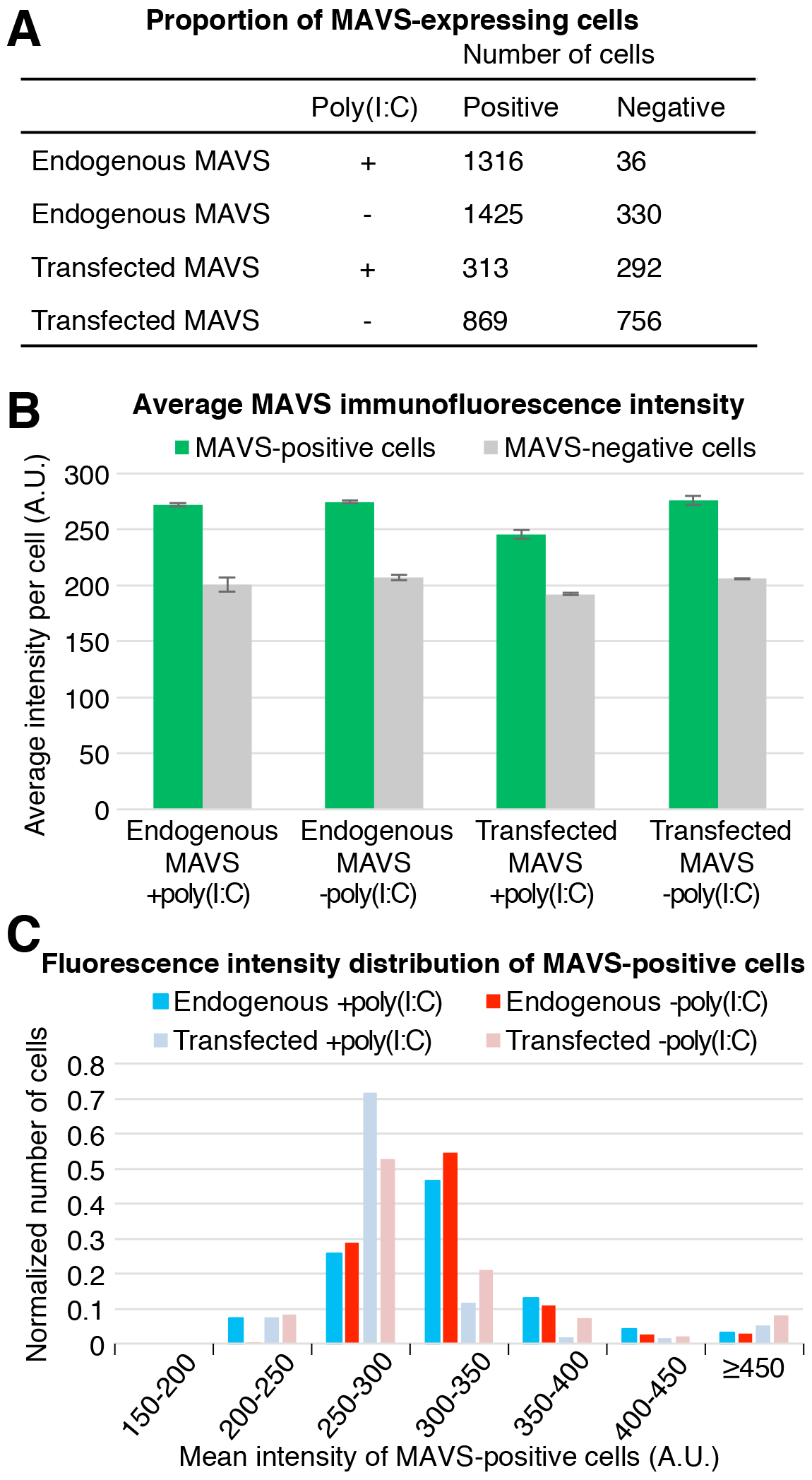
Comparison of MAVS immunofluorescence in wild-type MEFs and MAVS KO MEFs transfected with MAVS. (**A**) Numbers of MEFs expressing MAVS (positive) and lacking MAVS expression (negative) after (co)transfection with poly(I:C) (+poly(I:C)) or a control plasmid (-poly(I:C)). MAVS immunofluorescence was quantified with the General Analysis tool in Nikon Elements. A threshold of 215 arbitrary fluorescence intensity units (A.U.) was used as the cutofffor MAVS-positive cells. The transfection efficiency was approximately 50%. (**B**) Average MAVS immunofluorescence intensity of endogenous and transfected MAVS in MEFs (co)transfected with poly(I:C) (+poly(I:C)) or a control plasmid (-poly(I:C)). (**C**) The population distribution of the mean immunofluorescence intensity of MAVS-positive cells, plotted as the normalized number of cells versus fluorescence intensity from 150 to 450 A.U. in 50-A.U. bins.

### STORM shows MAVS signaling complexes are smaller than expected

Purified monomeric MAVS CARD spontaneously forms fibrils 0.2 - 1 μm in length [19, 21]. The fibrils, but not the monomers, activate IRF3 in cell-free assays [21]. However, SIM and STED imaging of cells with actively signaling MAVS did not resolve any clearly apparent fibrils (Fig. 1). To determine whether MAVS forms fibrils too small to resolve by SIM or STED, we employed a higher-resolution imaging modality, stochastic optical reconstruction microscopy (STORM), to image cells containing active MAVS signaling complexes. STORM can yield effective resolutions of 20 nm in the imaging plane [38]. MEFs were immunolabeled with an antibody against MAVS and a secondary antibody conjugated to Alexa Fluor 647, which was selected for its stable and prolonged signal in the STORM blinking buffer. MAVS KO MEFs were cotransfected with MAVS and poly(I:C) RNA as described for SIM and STED imaging. No MAVS filaments longer than the resolution limit were observed in the STORM images (Fig. 4). Instead, most of the MAVS immunofluorescence was present in fluorescent foci with irregular shapes, varying in diameter from 30 nm to 80 nm. These dimensions coincide with the effective resolution of the STORM imaging (see below). Unexpectedly, despite the global change in mitochondrial morphology associated with poly(I:C) treatment, there was no significant difference in the shapes and size range of MAVS foci in cells transfected with poly(I:C) or with a control plasmid.

**Fig. 4.**
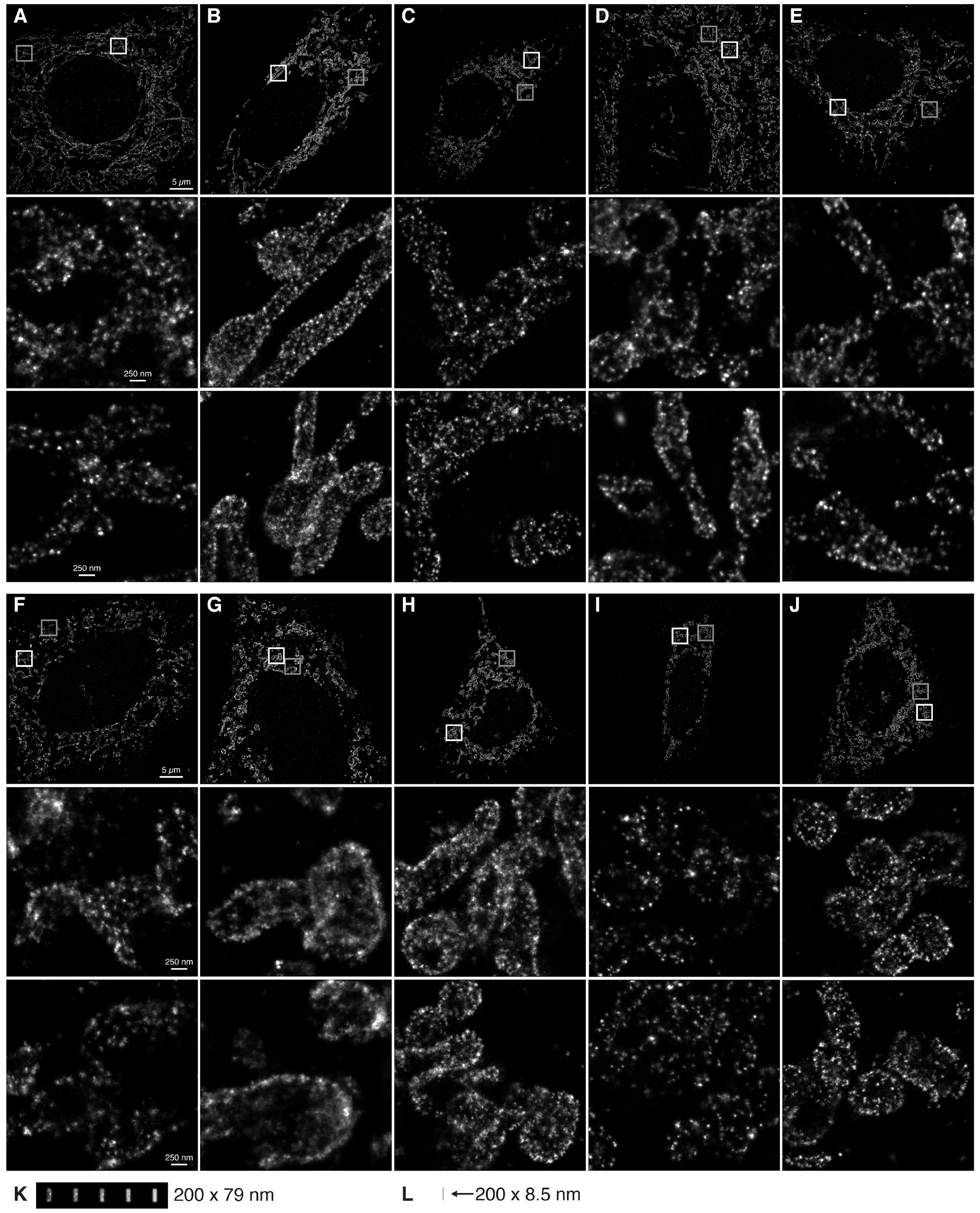
Stochastic optical reconstruction microscopy (STORM) of MAVS signaling complexes. (**A-E**) Five representative MAVS KO MEF cells transfected with MAVS and stained with an anti-MAVS antibody conjugated to Alexa Fluor 647. The boxed regions are shown enlarged in the lower panels of each overview panel (white box, middle panel; grey box, lower panel). (**F**-**J**) Five representative MAVS KO MEF cells cotransfected with MAVS and poly(I:C) RNA. The boxed regions are shown enlarged in the lower panels of each overview panel (white box, middle panel; grey box, lower panel). (**K**) Simulated renderings of immunofluorescence from MAVS fibrils with different antibody labeling efficiencies. An atomic model of a 200 nm MAVS CARD fibril was generated with one primary antibody and two secondary antibodies bound to each MAVS protomer (see Methods). The atoms of the lysine residues of each secondary antibody (the sites of fluorophore conjugation) were rendered with the same localization uncertainty as in the STORM images above. To emulate different antibody labeling efficiencies, the simulated immunolabeled fibrils were rendered with a given observation probability for each atom of 20%, 40%, 60%, 80% or 100% (from left to right in the panel). (**L**) A 200 nm MAVS CARD fibril rendered for reference on the same scale as the filaments in (**J**), without secondary antibodies and without blurring from localization uncertainty.

### Quantitative image analysis indicates MAVS fibrils are shorter than 80 nm

Since our ability to visualize submicrometer MAVS fibrils is critically dependent on the effective resolution of the imaging experiment, an accurate measurement of the imaging resolution is necessary to determine the minimum fibril length that can be resolved. We measured the resolution of our STORM images with the Fourier Ring Correlation (FRC) method, using an FRC of 0.143 as the threshold to measure resolution [39]. This criterion is the widely accepted standard for resolution assessment in cryo-electron microscopy (cryoEM) [40–42]. FRC curves calculated from our STORM images indicate that the resolution in the imaging plane ranged from 32 to 66 nm (Fig. 5A, B). Moreover, there was a correlation between the measured resolution of the images and the visual appearance of the MAVS foci. More specifically, images with the lowest resolutions (Fig. 4G, A) had more diffuse density, whereas images with the highest resolutions (Fig. 4E, I, C) had more pronounced foci, regardless of poly(I:C) treatment. This suggests that the appearance of clearly visible foci is determined by the imaging resolution rather than by the size of the imaged object. The calculated FRC resolution of the lowest-resolution STORM image was 66 nm. This is consistent with the SIM and STED data (Fig. 1A, C), which showed no correlation between poly(I:C) treatment and the appearance of clearly distinguishable MAVS fluorescence foci.

**Fig. 5.**
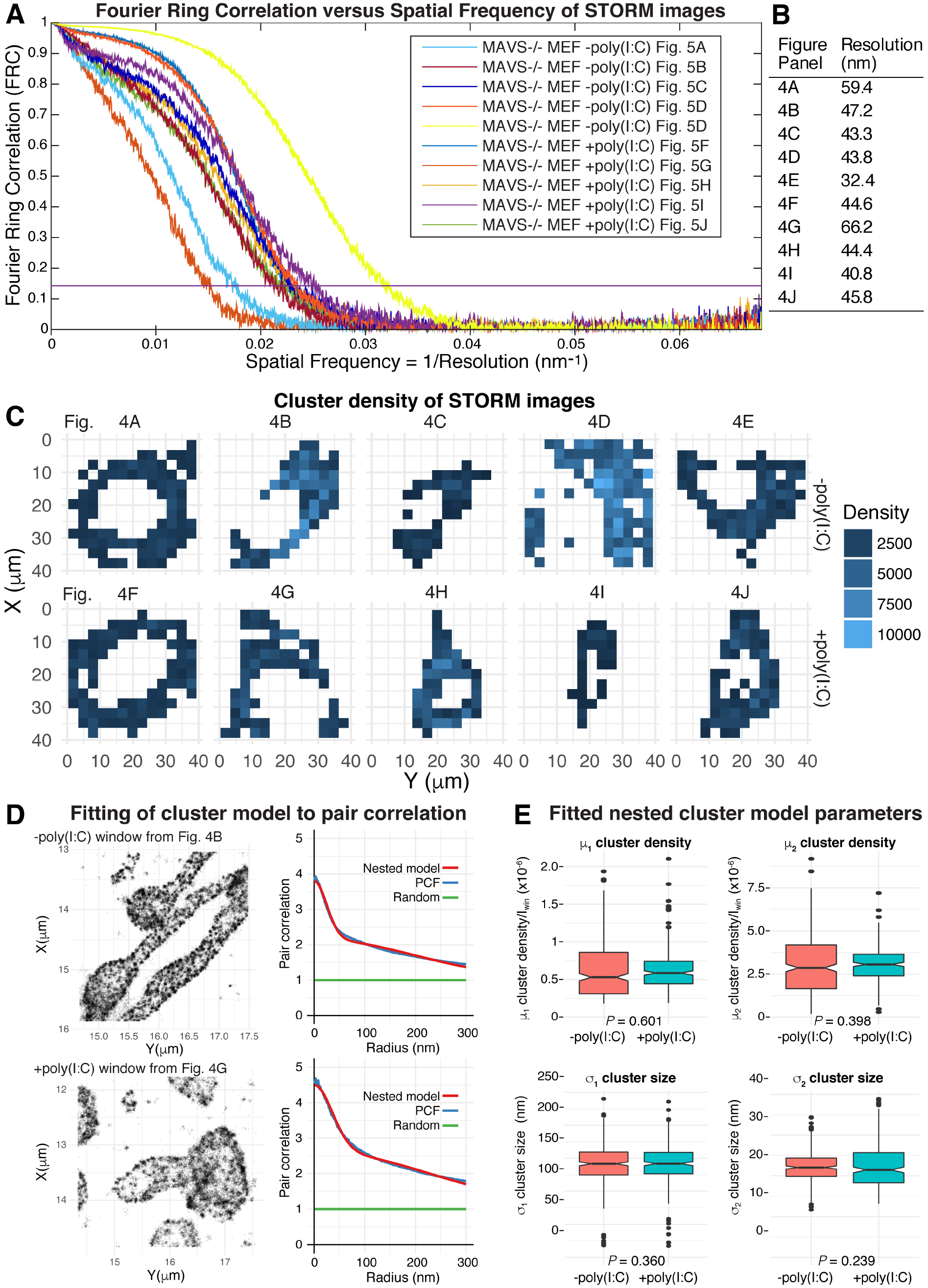
Resolution assessment and cluster analysis of STORM images. (**A**) Fourier Ring Correlation (FRC) curves were calculated for each STORM image shown in Fig. 4 (see Methods). An FRC value of 0.143, indicated by a horizontal purple line, was used as the threshold to measure resolution. (**B**) Effective resolution of each STORM image calculated from the FRC curves (FRC = 0.143). Cluster analysis of MAVS immunofluorescence in the STORM images. (**C**) Each image was divided into tiled 3 x 3 μm windows covering the field of view. Windows with a mean fluorescence intensity greater than half the mean intensity of the whole image were selected for analysis (shaded in blue). (**D**) A nested two-cluster model (red curve) fitted to the pair correlation function (PCF, blue curve), shown for representative individual cluster analysis windows from images of poly(I:C)-treated and control samples. (**E**) Box plots of the four parameters fitted in the cluster analysis, μ1, σ1, μ2, σ2, as defined in the Materials and Methods. The cluster density parameters, μ1 and μ2, were normalized against the mean intensity of the cluster analysis window, *I*_win_.

To quantify any subtle effects that poly(I:C) treatment might have on the clustering of MAVS foci, we performed cluster analysis on the STORM images using the pair correlation function. A hierarchical cluster model of two nested clusters was necessary to obtain a good fit to the pair correlation curve (Fig. 5C, D) [43, 44]. The clustering analysis showed that the two cluster size parameters of the nested cluster model were not significantly different in poly(I:C)-treated and control samples (Fig. 5E), confirming the lack of correlation between poly(I:C) treatment and the size of MAVS fluorescent foci.

Reconstruction of fine structural features by super-resolution microscopy depends on the precision with which the position of each fluorophore can be localized, and on the density and spatial distribution of active fluorophores along the labeled sample [39]. In the case of immunolabeled samples, the primary and secondary antibodies increase the spacing between the molecule of interest and the fluorophore significantly (by 20 - 35 nm) [45]. To determine whether 200 nm MAVS fibrils would in principle be visible in STORM images with the localization precision, labeling strategy and fluorophore properties inherent to our study, we generated an atomic model of 200 nm MAVS CARD fibrils bound to primary and secondary antibodies, and simulated the appearance of the fibrils with the same localization precision as our STORM images at different labeling efficiencies (Fig. 4K, L). Immunolabeling increases the overall diameter of the fibrils from 8.5 nm to 79 nm, but fibrils were nevertheless clearly visible with labeling efficiencies greater than 40%. Notably, the diameter of immunolabeled MAVS CARD fibrils is similar to the lowest resolution measured for the STORM images (79 vs. 66 nm, respectively). Therefore, immunolabeled MAVS fibrils shorter than 80 nm in axial length would be expected to appear as globular foci in STORM. Hence the absence of visible fibrils in our images remains consistent with the presence of helical MAVS fibrils up to 70 - 80 nm. A MAVS CARD helical assembly of this length would contain 136 - 156 MAVS molecules based on the 5.13 Å axial rise per protomer [19]. By comparison, purified MAVS fragments form filaments 200 - 1,000 nm in length in solution [19, 21].

### MAVS TM is required for mitochondrial remodeling and IFN-β signaling

MAVS CARD fibrils are sufficient to activate IRF3 in cytosolic extracts, but the transmembrane domain (TM) of MAVS is absolutely required for MAVS to activate IRF3 and induce interferon [5, 21]. We have shown that MAVS signaling activation causes changes in overall mitochondrial morphology similar to those associated with apoptosis, consistent with the documented proapoptotic activity of MAVS, which is dependent on the TM but not the CARD of MAVS [24]. To determine whether the MAVS TM is required for MAVS-dependent mitochondrial remodeling we used STED microscopy to image MAVS KO MEFs transfected with a plasmid encoding MAVS with the TM deleted (MAVS-ΔTM). As reported previously [5], we found that MAVS-ΔTM had diffuse cytosolic staining (Fig. 6A). Notably, cells expressing MAVS-ΔTM showed no visible aggregation, and little or no mitochondrial remodeling upon induction with poly(I:C), consistent with a recent report that the TM is required for the formation of high molecular weight MAVS aggregates [46].

**Fig. 6.**
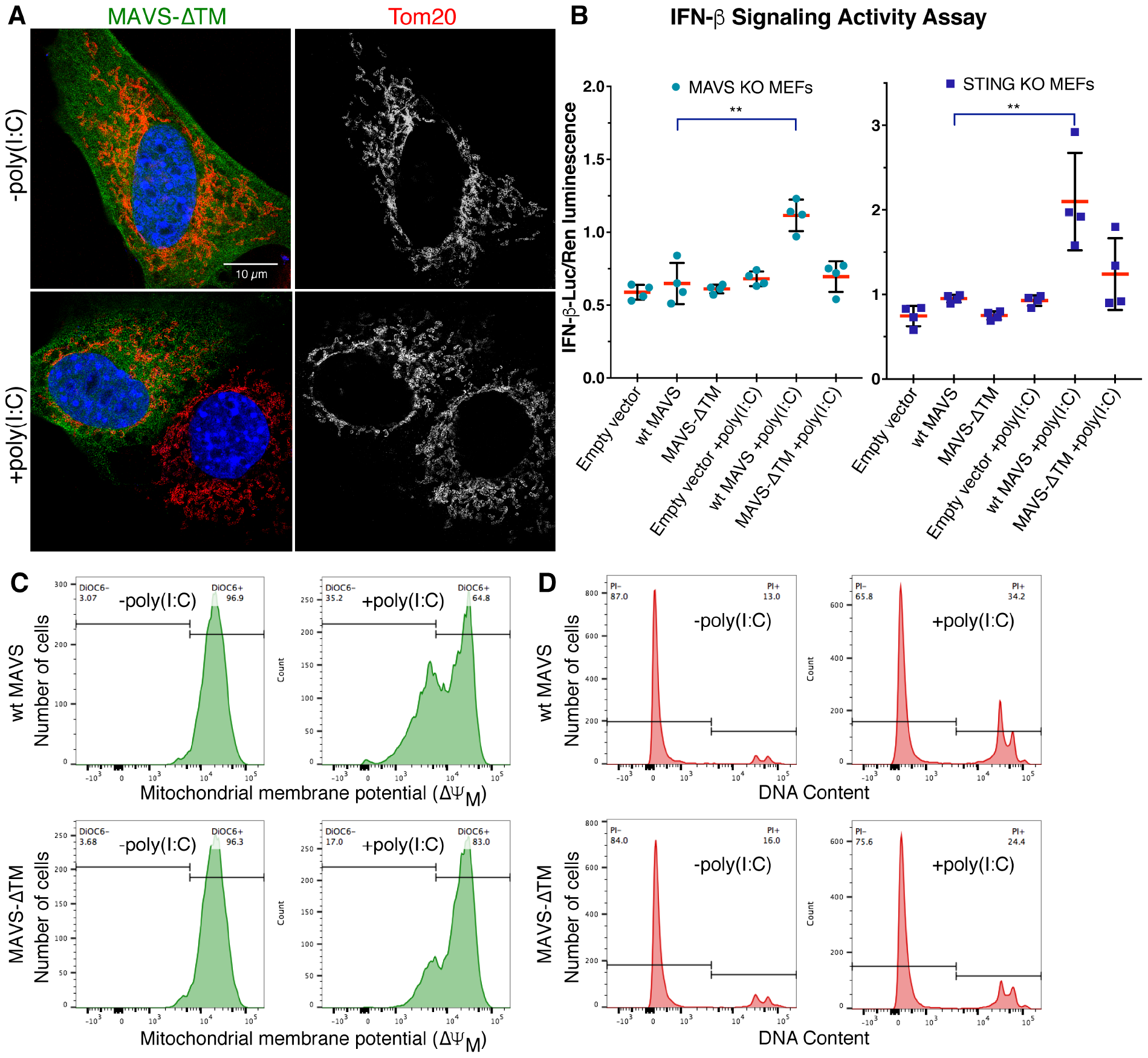
The MAVS transmembrane domain is required for mitochondrial remodeling, signaling and cell death activities of MAVS. (**A**) STED of MAVS-ΔTM and TOM20 in MAVS KO MEFs transfected with either MAVS-ΔTM and poly(I:C) RNA (+poly(I:C)) or with MAVS-ΔTM and a control plasmid (-poly(I:C)), with immunolabeling of MAVS-ΔTM (green), TOM20 (red) and DAPI staining (blue), or with TOM20 staining shown separately in grey. Only one of the two cells in the poly(I:C)-treated panel is positive for MAVS-ATM. (**B**) IFN-β dual-luciferase reporter assays in MAVS KO MEFs (left) and STING KO MEFs (right) co-transfected with plasmids encoding MAVS, poly(I:C) or control DNA, firefly luciferase under an IFN-β-inducible promoter and *Renilla* luciferase under a constitutive promoter. Relative luciferase activity was calculated as the ratio of firefly luciferase luminescence to *Renilla* luciferase luminescence. Error bars represent the standard deviation from the mean; n = 4. (**C**) Flow cytometry of DiOC6-stained MAVS KO MEFs cotransfected with wild-type MAVS or MAVS-ATM and poly(I:C) or a control plasmid. 35% of cells transfected with poly(I:C) and wild-type MAVS had a loss of inner mitochondrial membrane potential 16 h post-transfection, versus 17% of cells transfected with poly(I:C) and MAVS-ATM, and 3-4% of cells transfected with a control plasmid instead of poly(I:C). (**D**) Flow cytometry of PI-stained MAVS KO MEFs cotransfected with wild-type MAVS or MAVS-ΔTM and poly(I:C) or a control plasmid. 34% of cells transfected with poly(I:C) and wild-type MAVS had reduced nuclear DNA content 16 h post-transfection, versus 24% of cells transfected with poly(I:C) and MAVS-ΔTM, and 13-16% of cells transfected with a control plasmid instead of poly(I:C).

Previous studies have measured MAVS signaling activity from cytosolic or mitochondrial cell extracts. We confirmed that MAVS KO MEFs transfected with wild-type MAVS and poly(I:C) following the same protocol used for super-resolution imaging induced IFN-β signaling in the dual-luciferase reporter assay (Fig. 6B). In contrast, cells expressing MAVS-ATM failed to activate IFN-β signaling. The signal-to-noise ratio was low in the assay, however, due at least in part to induction of IFN-β signaling by cytosolic DNA sensing pathways such as cGAS-STING [47] in response to the transfected plasmid DNA. We therefore performed the luciferase reporter assay in STING KO MEFs (Fig. 6B), which are defective for cGAS-dependent DNA sensing [48]. The signal-to-noise was higher with STING KO MEFs than with MAVS KO MEFs despite the presence of endogenous MAVS in the STING KO MEFs. A slight but statistically insignificant increase in signaling was observed in STING KO MEFs transfected with MAVS-ATM. This is consistent with previous work showing that purified recombinant MAVS-ΔTM can, in its aggregated form, induce aggregation of endogenous wild-type MAVS and IRF3 activation in cell extracts enriched for mitochondria [21].

### MAVS induces cell death in response to cytosolic RNA

An early hallmark of apoptosis is the depolarization of the inner mitochondrial membrane [49], which is followed at later stages of cell death by loss of nuclear DNA content due to DNA fragmentation [50]. Overexpression of MAVS in HEK293T cells was shown previously to induce apoptosis [24]. To determine whether MAVS KO MEFs expressing physiological levels of MAVS induced apoptosis in response to activation with cytosolic dsRNA, we conducted cell death assays on cells transfected with MAVS and poly(I:C) RNA following the same protocol as for super-resolution imaging. Inner mitochondrial membrane depolarization and loss of nuclear DNA content were measured with the fluorescent markers 3,3'-dihexyloxacarbocanine iodide (DiOC6) (Fig. 6C) and propidium iodide (PI) (Fig. 6D), respectively. We found poly(I:C) treatment induced a loss of mitochondrial membrane potential in one third of cells transfected with wild-type MAVS, and loss of nuclear DNA content indicative of cell death in 21% of cells, at 16 h posttransfection, the same time point used for super-resolution imaging (Fig. 6C, D). In contrast poly(I:C) treatment of cells transfected with MAVS-ΔTM induced a loss of mitochondrial membrane potential and DNA content in only 13% and 8% of cells, respectively.

## Discussion

### Implications for signal transduction by MAVS

Taken together, our super-resolution light microscopy data suggest that in live cells, MAVS signaling complexes contain significantly smaller fibrils than those formed in vitro by soluble MAVS fragments. The absence of visible fibrils in SIM, STED and STORM images of cells expressing physiological levels of MAVS activated with various stimuli (synthetic or viral dsRNA), and a quantitative assessment of the resolution of the STORM images are all consistent with an upper limit in the range of 70 - 80 nm for the longest dimension of any MAVS fibrils within the MAVS signalosome. Unexpectedly, despite the global change in mitochondrial morphology associated with poly(I:C) treatment, there were no significant differences in the shape and distribution of MAVS foci in cells transfected with poly(I:C) or with a control plasmid. Cluster analysis of STORM images revealed no significant correlation between poly(I:C) treatment and the size of the clusters. Our confocal, SIM, STED and STORM data are consistent with each other and broadly consistent with previous fluorescence microscopy studies [21, 51], although a 3-D SIM reconstruction in one previous study showed MAVS forming rod-shaped puncta ranging from 100 nm to 650 nm in length, with a median length of 350 nm (n = 74) [27]. However, 350 nm is similar to the axial resolution of SIM and hence could be an overestimate of the MAVS cluster length.

An 80-nm long MAVS CARD helical assembly would still contain 156 MAVS molecules, a sufficiently large number to retain potential for polymerization-dependent signal amplification and to function as a mitochondrial signaling platform capable of recruiting the necessary number and diversity of downstream signaling proteins to elicit a robust IFN-β response [19, 20]. Given the smaller than expected size of MAVS signaling assemblies, a maximal MAVS-dependent signaling response may require a larger number of fibril nucleation events than previously thought. However, these nucleation events need not be independent, and MAVS signaling complexes may nevertheless assemble cooperatively, with assembled complexes promoting the nucleation of additional complexes without MAVS CARD filament extension beyond approximately 80 nm, but potentially forming two- or three-dimensional networks of microfibrils. Such a trade-off of reduced filament length in favor of increased nucleation was recently shown to occur elsewhere in the dsRNA sensing pathway. Indeed, LGP2 increases the initial rate of MDA5-dsRNA binding and limits MDA5 filament assembly, resulting in the formation of more numerous, shorter MDA5 filaments to generate a greater signaling activity [14]. Imaging at higher resolution, for example by electron microscopy, is required to elucidate the structural organization of MAVS signaling complexes.

We have shown that the MAVS transmembrane anchor is required for MAVS-associated mitochondrial remodeling and confirmed that the TM is required for MAVS-dependent IFN-β signaling and apoptosis in live cells and under the same experimental conditions used for the super-resolution imaging experiments in this study. The diffuse cytosolic localization of MAVS-ATM suggests that MAVS oligomerization is regulated by elements between the CARD and TM, and that the TM is required to overcome this regulation. It is tempting to speculate that the physical constraints imposed by MAVS aggregation and association with RLR-RNA complexes exert a physical force on the outer mitochondrial membrane capable of inducing distortions that could conceivably contribute to mitochondrial remodeling or leakage associated with apoptotic cell death.

## Materials and methods

### Expression plasmids

Genes encoding full-length mouse MAVS (residues 1-503, Uniprot entry Q8VCF0) and mouse MAVS with the transmembrane anchor (residues 479-503) deleted (MAVS-ATM) were cloned into the pCMV-SPORT6 vector (BioCat, Heidelberg, Germany).

### Cell culture

Wild-type mouse embryonic fibroblasts (MEFs), MAVS knockout (MAVS KO) MEFs, STING KO MEFs and 3T3 cells were cultured in Dulbecco’s modified Eagle Medium high glucose (DMEM) (Gibco, Waltham, MA), supplemented with 10% v/v fetal bovine serum and 10% sodium pyruvate (Gibco). Cells were maintained at 37°C in a 5% CO_2_ atmosphere.

### Transfections

Transient transfections were performed with the Neon transfection system (ThermoFisher, Waltham, MA) following the manufacturer’s protocol. For transfections in 6-well plates, the cells were transfected at a density of 2.5 x 10^5^ cells per well using a 10 μl Neon Tip and a total of 0.5 μg of nucleic acids per well. For transfections in 96-well plates, cells were transfected and seeded at 8.5 x 10^4^ cells per well using a 10 μl Neon Tip and a total of 0.1 μg of nucleic acids per well. For MAVS KO MEFs transfected with MAVS the transfected nucleic acids consisted of 1:1 (w/w) pCMV-MAVS plasmid DNA:poly(I:C) RNA (Midland Certified Reagents), or 1:1 (w/w) pCMV-MAVS plasmid DNA:pCMV empty vector plasmid DNA for the untreated control. The total amount of poly(I:C) RNA was 0.6 - 1 ng per 1,000 cells. For the MAVS KO MEF negative control and the experiments with wild-type MEFs and endogenous MAVS (Fig. 1B, 3) the transfected nucleic acids consisted of 1:1 (w/w) pCMV empty vector plasmid DNA:poly(I:C) RNA, or pCMV empty vector plasmid DNA only for the untreated control.

### West Nile Virus infection

West Nile reporter virus particles were generated in HEK 293T cells by cotransfecting the cells with plasmids encoding the structural proteins (C, prM, E) and a subgenomic replicon containing GFP (pWNVII-Rep-G-Z), as described previously [52, 53]. The plasmids were kind gifts from Theodore Pierson (NIH). Supernatant collected 48 h after transfection was filtered through 0.2 μm membranes and used to infect 3T3 cells. 24 h postinfection, the cells were fixed with 4% formaldehyde solution and labeled for immunofluorescence.

### Immunofluorescence staining

For anti-MAVS staining cells were fixed 16 h posttransfection on the microscopy coverslips with 4% formaldehyde in phosphate-buffered saline (PBS), permeabilized in 50 mM Tris/HCl pH 7.5, 0.15 M NaCl, 0.02% Saponin (TBSS) for 30 min at 37°C, and then blocked in 5% BSA in TBSS for 1 h at 37°C. Cells were stained with a mouse anti-MAVS monoclonal IgG2a antibody (MAVS (C-1), Santa Cruz Biotechnology, Dallas, TX) in TBSS at 1:80 dilution for 1.5 h at 37°C. After four 5-min washes with TBSS, the cells were incubated with goat anti-mouse IgG conjugated to Alexa Fluor 488 (Molecular Probes, Eugene, OR) at 1:80 dilution for 1 h at room temperature, washed with TBSS, and mounted for imaging with ProLong Gold Antifade Mountant with DAPI (ThermoFisher). For anti-TOM20 staining cells were fixed and blocked the same way as for anti-MAVS staining. TOM20 was stained with rabbit anti-TOM20 polyclonal IgG antibody (TOM20 (FL-145), Santa Cruz Biotechnology) in TBSS at 1:80 dilution for 1.5 h at 37°C. After washing with TBSS, the cells were incubated with goat anti-rabbit IgG conjugated to Alexa Fluor 555 (Molecular Probes) at 1:80 dilution for 1 h at room temperature, washed with TBSS, and mounted for imaging with ProLong Gold Antifade Mountant with DAPI (ThermoFisher). For the MAVS costaining with IRF-3 and MDA5 cells were fixed, permeabilized, and blocked as for anti-MAVS staining above. IRF-3 and MDA5 were stained with rabbit anti-IRF3 polyclonal IgG (IRF-3 (FL-425), Santa Cruz Biotechnology) and rabbit anti-MDA5 polyclonal IgG (Enzo Life Sciences, Farmingdale, NY), respectively, at 1:80 dilution in TBSS for 1.5 h at 37°C. After washing with TBSS, the cells were incubated with goat anti-rabbit antibody conjugated to Alexa Fluor 647 (Molecular Probes) at 1:80 dilution for 1 h at room temperature, and then mounted for imaging with ProLong Gold Antifade Mountant with DAPI (ThermoFisher).

### Mitochondrial remodeling analysis

Widefield images were acquired at 100x magnification with an inverted Nikon TE2000 microscope equipped with a Niji LED illumination (Bluebox Optics, Huntingdon, UK), a 100x/1.49NA oil objective (Nikon, Tokyo, Japan) and NEO sCMOS camera (Andor, Belfast, UK). A custom ImageJ macro script (Mitochondria_remodelling_analysis.ijm in Supplementary Data) was written to analyze mitochondrial distance, area and length parameters with ImageJ [54]. More specifically, distance from the nucleus was measured as the shortest distance from each TOM20 fluorescence pixel to a DAPI fluorescence pixel. Second, the fraction of the cytosolic area occupied by mitochondria was measured in each cell. Third, the length of mitochondrial compartments was measured by skeletonizing the mitochondrial fluorescence signal, segmenting the skeleton at branchpoints and measuring the length of each resulting segment. These measurements were performed on 28 cells without poly(I:C) and 11 cells with poly(I:C) treatment.

### MAVS-MDA5 interaction analysis

The averaged distances of the collective points in the MDA5 and MAVS fluorescence signals after treatment with poly(I:C) or a control plasmid was calculated with the Interaction analysis plugin for ImageJ from MosaicSuite (MOSAIC Group, Dresden, Germany) [35]. Colocalization was measured by calculating the Pearson correlation coefficient with the Coloc 2 plugin in ImageJ.

### Confocal fluorescence imaging

Confocal imaging (Figs. 1B, 1D, 2, 3) was carried out on a Zeiss (Oberkochen, Germany) LSM 780 or Leica (Wetzlar, Germany) TCS SP8 laser scanning inverted microscope with a 40x/1.3 NA or 63x/1.4 NA oil immersion objective lens. The Zeiss was equipped with diode 405 nm, Argon multiline, DPSS 561 nm and HeNe 633 nm laser lines.

The Leica was equipped with diode 405 nm and NKT Super K pulsed white light (470-670 nm) lasers. On the Zeiss, DAPI/Alexa Fluor 405 (405 nm excitation), Fluorescein/ATTO488 (488 nm excitation), mApple (561 nm excitation), Alexa Fluor 647/SiR-647 (633 nm excitation) and differential interference contrast (DIC) images were collected and analyzed with ZEN 2009-2011 (Zeiss) and ImageJ [54]. On the Leica, Alexa Fluor 405 (405 nm excitation), GFP/ATTO488 (488 nm excitation) and Alexa Fluor 647 (633 nm excitation) images were collected and analyzed with LAS AF (Leica). Images were deconvolved with Huygens Professional (Scientific Volume Imaging, b.v., Hilversum, The Netherlands).

### Structured Illumination Microscopy (SIM)

Cells were plated as a monolayer on high performance Zeiss cover glasses (D = 0.17 mm, 18 x 18 mm type 1½ H, ISO 8255-1 with restricted thickness-related tolerance of ± 0.005 mm, refractive index = 1.5255 ± 0.0015) in 6-well CytoOne TC-Treated tissue culture plates (STARLAB, Milton Keynes, UK). Images were acquired on a Zeiss ELYRA S.1 system with a pco.edge 5.5 scientific CMOS camera using a 63x/1.4 NA oil immersion objective lens. For optimal lateral spatial sampling a 1.6x intermediate magnification lens was used, resulting in a pixel size of 64 nm. Excitation was achieved with a 488 nm laser and emission was filtered with a 495-550 nm band pass filter. Modulation of the illumination light was achieved using a physical grating with 28 μm spacing. This patterned illumination was shifted through 5 phases at each of 3 rotational angles per image. Raw data were processed using ZEN software.

### STimulated Emission Depletion (STED) microscopy

Cells were plated as a monolayer on high performance Zeiss cover glasses (D = 0.17 mm, 18 x 18 mm type 1½ H, ISO 8255-1 with restricted thickness-related tolerance of ± 0.005 mm, refractive index = 1.5255 ± 0.0015) in 6-well CytoOne TC-Treated tissue culture plates (STARLAB). Images were acquired on a Leica TCS SP8 STED confocal system with time-gated HyD GaAsP detectors using a 100x/1.4 NA oil immersion objective lens. Excitation was achieved with a NKT Super K pulsed white light (470 - 670 nm) laser tuned to 488 nm and STED was induced with a 592 nm laser (~40 MW cm^-2^). A time gate window of 2 - 6 ns was used to maximize STED resolution. 20 nm pixels were used to ensure adequate spatial sampling to support maximal expected resolution. Images were deconvolved with Huygens Professional.

### Stochastic Optical Reconstruction Microscopy (STORM)

In STORM, fluorophores are induced to switch, or “blink” between fluorescent and dark states. With a sufficiently small fraction of fluorophores in the fluorescent state at any given time, spatial overlap between adjacent fluorophores is avoided and the positions of individual fluorophores can be determined with high precision using the point-spread function of each fluorophore [38]. STORM images are generated by plotting the positions of each fluorophore blinking event as points in the image plane and applying to each point a blurring factor corresponding to the localization precision for that point. The appearance of the images also depends on the efficiency of fluorescent labeling and the extent of fluorophore bleaching during the experiment.

Cells were plated as a monolayer on high performance CellPath (Newtown, UK) HiQA Coverslips (No. 1.5 H, 24 mm Ø) in 6-well CytoOne TC-Treated tissue culture plates. Alexa Fluor 647-labeled samples were imaged in the following buffer condition: 10% w/v glucose, 0. 5 mg ml^-1^ glucose oxidase (Sigma-Aldrich, St. Louis, MO), 40 μg ml^-1^ catalase (Sigma Aldrich), and 0.1 M β-mercaptoethylamine (MEA) (Sigma-Aldrich) in PBS, pH adjusted to 7.4 with concentrated HCl. To minimize air oxidation of the fluorophore, the imaging well was filled to full capacity with the imaging buffer and sealed with a cover glass. Imaging was performed with an N-STORM microscope (Nikon) equipped with an Apochromat TIRF 100x/1.49 NA oil immersion objective lens, a single photon detection iXon Ultra DU897 EMCCD camera (Andor), and 405 nm (30 mW) and 647 nm (170 mW) laser lines. To acquire single-molecule localizations, samples were constantly illuminated with the 647 nm laser at 2-3 kW cm^-2^. To maintain adequate localization density, the 405 nm laser was used to reactivate the fluorophore. During imaging, Perfect Focus System was used to maintain the axial focal plane. 7 - 10 x 10^4^ frames were collected for each field of view at a frame rate of 50-70 Hz. STORM images were reconstructed with NIS Elements Advanced Research (Nikon): lateral drift correction was performed using automated cross-correlation between frame sets and each localization point in the reconstructed image was shown as a normalized Gaussian, the width of which corresponded to the localization uncertainty calculated using the Thompson-Larson-Webb equation [55]. Each STORM image contained 1-2 million molecules within a single cell.

### Fourier ring correlation (FRC)

The effective optical resolution of the STORM images was assessed using the FRC method proposed by Nieuwenhuizen *et al*. (2013) using the Matlab code provided [39] (frc_analysis.m in Supplementary Data). An FRC threshold of 1/7 (0.143) was used to determine the resolution of each rendered STORM image.

### Cluster analysis of MAVS immunofluorescence in STORM images

Cluster analysis was performed to assess the size of fluorescent foci from immunolabeled MAVS in the STORM images. Using a custom C++ script (nestedclusteranalysis.cpp in Supplementary Data), each image was divided into 3 x 3 μm windows covering the field of view and windows with a mean fluorescence intensity greater than an arbitrary threshold—half of the mean fluorescence intensity of the whole image—were selected for analysis (Fig. 5C-E). Cluster analysis was performed by fitted a hierarchical nested cluster model, or clusters of clusters model, to the pair correlation function (or radial distribution function), defined as the average number of points located in a ring of radius *r* centered around each point and normalized by the expected intensity taking into account the border of the window of analysis [44, 56]. The nested cluster model was fitted to the pair correlation curve using a least squares approach. The selected model comprised two clusters (smaller “inner” clusters that cluster into larger “outer” clusters) defined by four parameters (*μ*1, *σ*1, *μ*2, *σ*2) through the following equation:

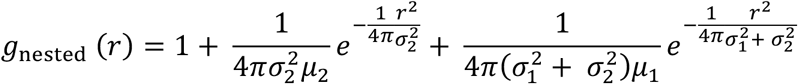

with σ1 and σ2 corresponding to the outer and inner cluster sizes, respectively.

### Simulated rendering of immunolabeled

MAVS fibrils. The atomic coordinates of an mouse antibody molecule (PDB code 1IGY) [57] were manually docked with UCSF Chimera [58] to an arbitrary epitope on the MAVS CARD coordinates (PDB code 3J6J) to simulate a primary antibody-MAVS complex. Two additional antibody molecules were then docked to two different arbitrary epitopes in the constant region (Fc domain) of the primary antibody. The helical symmetry of the MAVS CARD filament determined by cryo-electron microscopy (cryoEM) image reconstruction [19] was then applied consecutively to the atomic coordinate of the complex to generate a 200 nm MAVS CARD filament with bound primary and secondary antibodies. The simulated epitopes and orientations of the antibodies were selected to minimize steric clashes after the helical symmetry of the MAVS CARD filament was applied. A custom C++ code was used to render the atoms of the lysine residues, the sites of fluorophore conjugation, of each secondary antibody in the filament. Each atom was rendered on a two-dimensional image at a 5 nm scale with a given observation probability ranging from 20% to 100%, in order to emulate different antibody labeling efficiencies. The resulting image was then blurred in order to take into account the localization uncertainty of the STORM singlemolecule localizations events, approximately 10 nm.

### Dual-luciferase reporter cell signaling assay

MAVS KO MEFs or STING KO MEFs were transfected with wild-type MAVS or MAVS-ΔTM and poly(I:C) with the Neon transfection system as described above, using the same conditions as for light microscopy, except that plasmids encoding firefly luciferase under an interferon-β (IFN-β) promoter and *Renilla* luciferase under a constitutive promoter (Promega, Madison, WI) were also included in the transfection. Cells were transfected and seeded in quadruplet sets for each condition in 96-well clear bottom black polystyrene microplates (Corning, Corning, NY) at 5 - 6 x 10^3^ cells per well. Cells were lysed and the lysates transferred to all-white 96-well flat solid bottom plates suitable for tissue culture and luminescence measurements (Greiner Bio-One, Kremsmünster, Austria). Sample preparation from cell lysates and luciferase luminescence measurement were performed according to the manufacturer’s protocol for the Dual-Luciferase Reporter Assay System (Promega). IFN-β-dependent induction of firefly luciferase was measured in cell lysates 16 h post-induction with a PHERAstar (Ortenberg, Germany) FSX microplate reader with dual sample injection. IFN-β signaling activity was measured as the ratio of firefly luciferase luminescence to *Renilla* luciferase luminescence.

### Quantification of cell death

Depolarization of the inner mitochondrial membrane was quantified by staining cells with 40 nM of 3,3'-dihexyloxacarbocanine iodide (DiOC6, Sigma-Aldrich) or 6 mg ml^-1^ propidium iodide (PI) for 1.5 h in PBS. DiOC6 and PI fluorescence was quantified by flow cytometry. Quantification was set at 5,000 for cell count and FL1 (530 nm) and FL3 (670 nm) were used to acquire DiOC6 and PI signals, respectively. Data were acquired on a LSR II flow cytometer (BD Biosciences, San Jose, CA) and analyzed with FlowJo (TreeStar, Ashland, OR).

### Statistical analysis

No statistical methods were used to predetermine sample size, experiments were not randomized, and the investigators were not blinded to experimental outcomes. Unless otherwise noted, errors are presented as the standard deviation of the mean of four replicates conducted in a single independent experiment. Statistical significance was calculated with Prism 8 (GraphPad Software, San Diego, CA) using an unpaired t-test without prior assumptions regarding the standard deviations of each set of quadruplet measurement. Statistical significance was assigned as follows: *, P < 0.05; **, P < 0.01; ***, P < 0.001, n = 4.

## Author contributions

Conceptualization, M.-S.H. and Y.M.; Methodology, all authors; Software, J.B. and L.M.; Formal Analysis, J.B., M.P., L.M.; Investigation, M.-S.H., J.B., J.H., A.A., M.P.; Writing - Original Draft, M.-S.H. and Y.M.; Writing - Review & Editing, Y.M., with input from all authors; Visualization, M.-S.H., J.B., J.H., A.A., M.P. and Y.M.; Supervision, Y.M.; Project Administration, Y.M.; Funding Acquisition, Y.M.

## Acknowledgements

We thank Theodore Pierson (NIH) for his kind gift of plasmids encoding the West Nile reporter virus. We thank Michael Gale Jr. (Univ. of Washington) and Jan Rehwinkel (Univ. of Oxford) for their kind gifts of MAVS KO MEFs and STING KO MEFs, respectively. We thank Nick Barry and Ben Sutcliffe at the MRC-LMB Light Microscopy Facility for support with the collection and handling of light microscopy data. We thank Yangci Liu and Clare Bryant for their comments on the manuscript, and all members of the Modis lab for insightful discussions. This work was supported by a Wellcome Trust Senior Research Fellowship to Y.M. (101908/Z/13/Z).

